# Exploring the role of prior exposure and image quality in neural and behavioural prediction effects

**DOI:** 10.64898/2026.07.15.738617

**Authors:** Phuong N.U. Dang, Jason B. Mattingley, Margaret J. Moore

## Abstract

According to predictive processing theories, the brain achieves efficiency in perception by comparing current sensory inputs with stored representations of the external world. Previous research has shown that manipulations of expectation can alter neural processing of low-level visual features as well as more complex, naturalistic objects. It remains unclear, however, precisely how the effects of expectancy are altered through learning and changes in stimulus fidelity. Here, we characterised behaviour and neural activity patterns while systematically varying individuals’ prior exposure to object sequences and the quality of the stimuli therein. Participants viewed rapid image sequences while we recorded their brain activity using electroencephalography (EEG). The stimulus sequences were probabilistically structured such that the appearance of each object was either expected, unexpected, or random. Participants were faster and more accurate in detecting cued target stimuli when these were expected relative to unexpected or random. Multivariate analysis of EEG activity patterns time-locked to the appearance of object images revealed reliably reduced decoding accuracy for both expected and unexpected stimuli relative to random stimuli. These patterns were consistent across exposure and image quality conditions, though exploratory analyses suggested that subtle changes in neural prediction effects may be linked to exposure. The findings do not provide strong support that the brain differentially represents expected versus unexpected object information and suggest that exposure and image quality do not reliably interact with predictive processes in object recognition.

## Introduction

Prediction is proposed to be an organising principle of brain function supporting sensory processing, perception, and action (Clark, 2013; Friston, 2005, 2010; Lee & Mumford, 2003; Rao & Ballard, 1999; Summerfield & de Lange, 2014). According to predictive processing theory, incoming sensory inputs are differentially processed depending on their alignment with the brain’s computational expectations. These processes have been associated with distinct neural activity patterns for naturalistic objects that are expected versus unexpected (Kaposvari et al., 2016; Kumar et al., 2017; Moore et al., 2024, 2025; Richter et al., 2018, 2024). Naturalistic objects are uniquely valuable for exploring neural prediction effects as they require multi-level hierarchical visual processing and engage a wider network of brain areas than elementary visual features such as orientation, spatial frequency and colour (Felleman & Van Essen, 1991; Friston, 2005; Grill-Spector, 2013; Grill-Spector & Malach, 2004; Groen et al., 2017; Lee & Mumford, 2003; Ungerleider & Haxby, 1994).

Within experimental settings, expectations are often learned through exposure to probabilistic information (de Lange et al., 2018; Ferrari et al., 2022; Knill, 2007). Often, however, such predictive relationships take time to be learned, either consciously or unconsciously (Bays et al., 2015; Ferrari et al., 2022; Turk-Browne et al., 2005), implying that expectancy effects on neural representations may evolve with continued exposure. However, previous neural studies have varied widely in the degree of exposure used (Kumar et al., 2017; Richter et al., 2018, 2024; Moore et al., 2024). The impact of exposure differences on neural responses to expectation has not been explicitly investigated, and addressing this would illustrate how prediction effects develop with experience.

Under natural viewing conditions, visual objects are often degraded or occluded, which can lead to perceptual errors and corresponding reductions in the fidelity of associated neural representations (Erlikhman & Caplovitz, 2017; Grootswagers et al., 2019; Li et al., 2026; Moore et al., 2024). Behavioural experiments have indicated that prediction exerts its strongest effects on perception when image quality is degraded (Brandman & Peelen, 2017; Carrez-Corral et al., 2025; Rossel et al., 2023; Stein & Peelen, 2015). In addition, Moore et al. (2024) recently found a larger boost in neural decoding of degraded expected visual stimuli relative to intact stimuli. This result suggests that fulfilled predictions may facilitate veridical perception, but it is unclear whether improved representational fidelity corresponds to improved behavioural performance. Determining how image quality influences both neural and behavioural prediction effects would provide a more comprehensive understanding of the impact of prediction on visual object recognition.

Visual processing is thought to depend on both prediction strength and stimulus quality (de Lange et al., 2018; Knill & Pouget, 2004). Prediction strength can be operationalised as the extent to which participants have been exposed to relevant probabilistic information, while stimulus quality can be manipulated through varying levels of image degradation (Friston 2005; de Lange et al., 2018). Prediction strength and stimulus quality have been proposed to interact, such that stronger predictions have a greater influence on visual processing when the sensory input is expected but degraded (Brogaard & Sørensen, 2023; Kersten et al., 2004). Previous studies using elementary stimulus features such as orientations and dots have supported this proposal (Goodwin et al., 2025; Tomassini et al., 2010; Vilares et al., 2012). However, it is not clear how prior exposure and image quality jointly modulate predictive effects for naturalistic object stimuli.

Here, we characterised how prior exposure and image quality influence prediction effects by measuring behavioural and electroencephalography (EEG) responses to rapid probabilistic sequences of intact and degraded object stimuli across two experimental sessions. If stronger predictions emerge with prior exposure, prediction effects should be more pronounced in the second testing session relative to the first. If prediction effects are modulated by image quality, the magnitude of any prediction effects should be greater in degraded relative to intact stimuli. Finally, if prior exposure strengthens predictions which then interact with image quality, prediction effects should be most pronounced in degraded stimuli during the second testing session.

## Methods

### Participants

Forty-four participants were recruited through The University of Queensland (UQ) and were compensated at a rate of 20 AUD per hour. Participants completed two testing sessions, one involving EEG and the other behavioural only (order counterbalanced; *M* = 3.55 days apart, *SD* = 3.26; time ≈ 2 hours per session). Four participants were excluded because they did not complete both testing sessions. Data from 40 participants (35 females; 5% left-handed; *M* = 24.25 years old, *SD* = 4.78, range = 19–39) were analysed. All participants had normal or corrected-to-normal vision (*M* = 8.30 Snellen score, *SD* = 1.2). The study was approved by UQ’s Human Research Ethics Committee (HREA: 2024/HE001654).

### Stimuli, Apparatus and Design

EEG and behavioural responses were recorded as participants viewed rapid serial visual presentation (RSVP) sequences of intact and degraded real-world object stimuli, while providing button press responses to specific stimuli.

Stimuli included nine naturalistically coloured objects obtained from an online image database (www.pngimg.com; Fig.1a). Similar stimuli have yielded reliable decoding of object-specific information in previous RSVP paradigms (Grootswagers et al., 2019; Moore et al., 2024). To generate degraded stimuli, diffeomorphic warping (Stojanoski & Cusack, 2014) was applied (10^th^ step in a 7-step circular warping matrix, maximum distortion level = 20). The warping intensity was selected in line with the recommendations of Stojanoski and Cusack (2014) to diminish, but not eliminate, recognisability. Using these parameters, three different degraded versions of each stimulus were generated using the same level of distortion but different warping matrices (Fig.1b).

**Figure 1:**
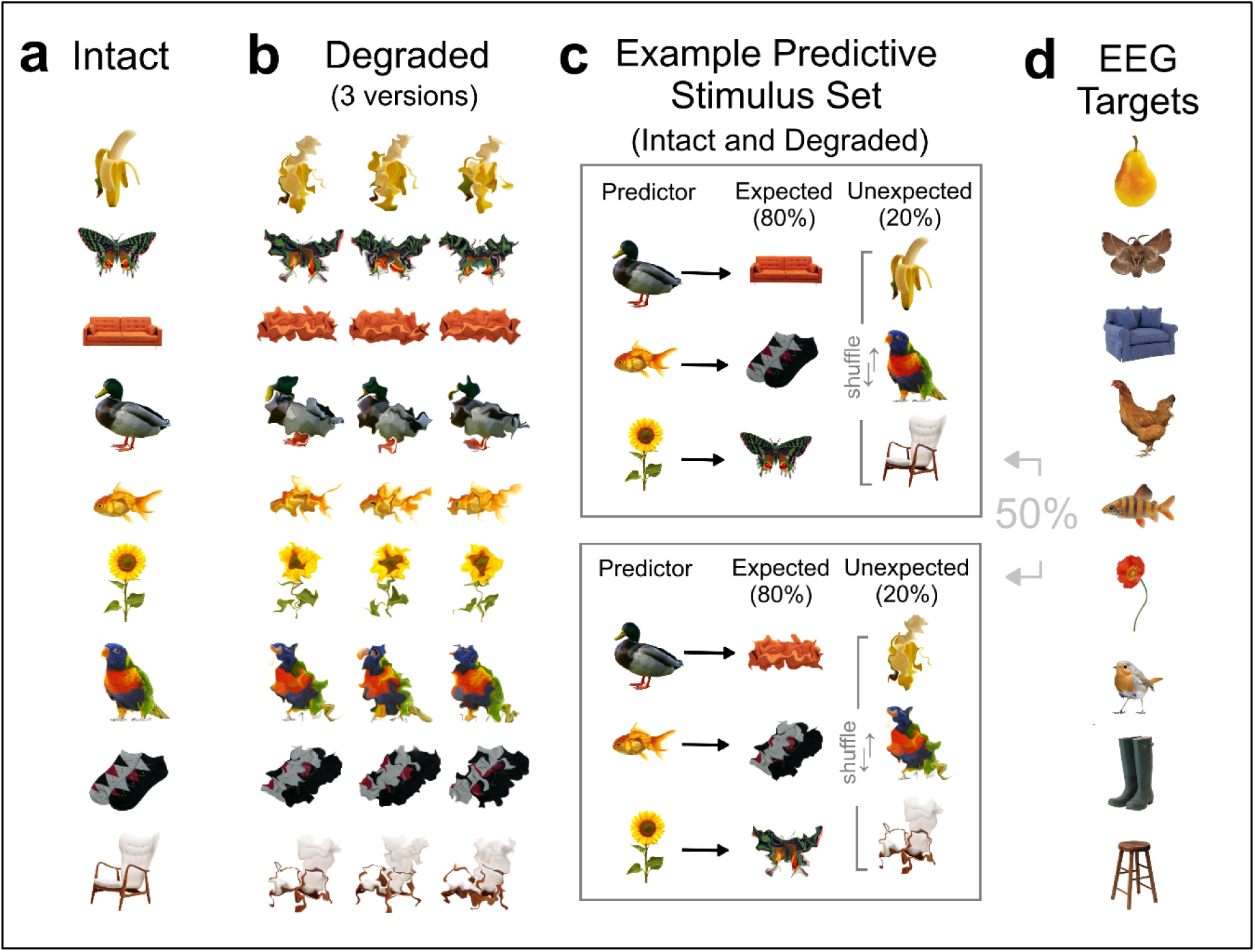
Visual object stimuli used in the experiment. **(a)** Intact stimuli used to construct predictive stimulus sets. **(b)** Degraded stimuli with three different warping matrices. Degraded stimuli for the EEG experiment only included the left-most column of warped objects. **(c)** An example stimulus set with predictive relationships. Half of expected and unexpected stimuli are intact (top panel), and half are degraded (bottom panel). **(d)** Target stimuli (‘attention probes’) for the EEG task were from a similar category to those used in the main stimulus sets.

Perceptual expectations were induced using a probabilistic transition structure. ‘Predictor’ objects (*n* = 3) were followed by a specific ‘expected’ object in 80% of trials. In the remaining 20%, one of three possible ‘unexpected’ objects was presented instead (Fig.1c). To equalise stimulus presentation frequency, unexpected objects could also appear in sequence positions that did not follow a predictor stimulus (i.e., randomly). Half of all objects in expected, unexpected and random sequence positions were intact, and half were degraded. To control for any subtle differences in the distribution of low-level object features (e.g., colour, orientation), for each participant, one of four possible stimulus sets was used in which different objects acted as predictors, expected, and unexpected stimuli (Fig.S1). Participants were not informed about the predictive relationships. We also included a separate set of objects as ‘attention probe’ targets, which participants responded to during EEG recordings (Fig.1d).

Stimuli were presented in 5 Hz RSVP sequences (100 ms display time, 100 ms interstimulus blank). Different monitors were used in the EEG (100 Hz ViewPix3D) and behavioural (120 Hz Asus VG248QE) sessions. In all sessions, stimuli were presented centrally, subtending 3° visual angle, against a white background at an approximate viewing distance of 60 cm. Participants were instructed to maintain fixation and make responses with their right hand using the down-arrow button on a standard computer keyboard.

### Behavioural Paradigm

The behavioural paradigm evaluated whether expectancy modulates behavioural performance (speed and proportion correct). Participants viewed 240 probabilistically structured sequences of intact and degraded stimuli (∼2400 displays per predictive pair). Each sequence contained 10 displays per stimulus (intact and degraded inclusive; Fig.2a).

**Figure 2:**
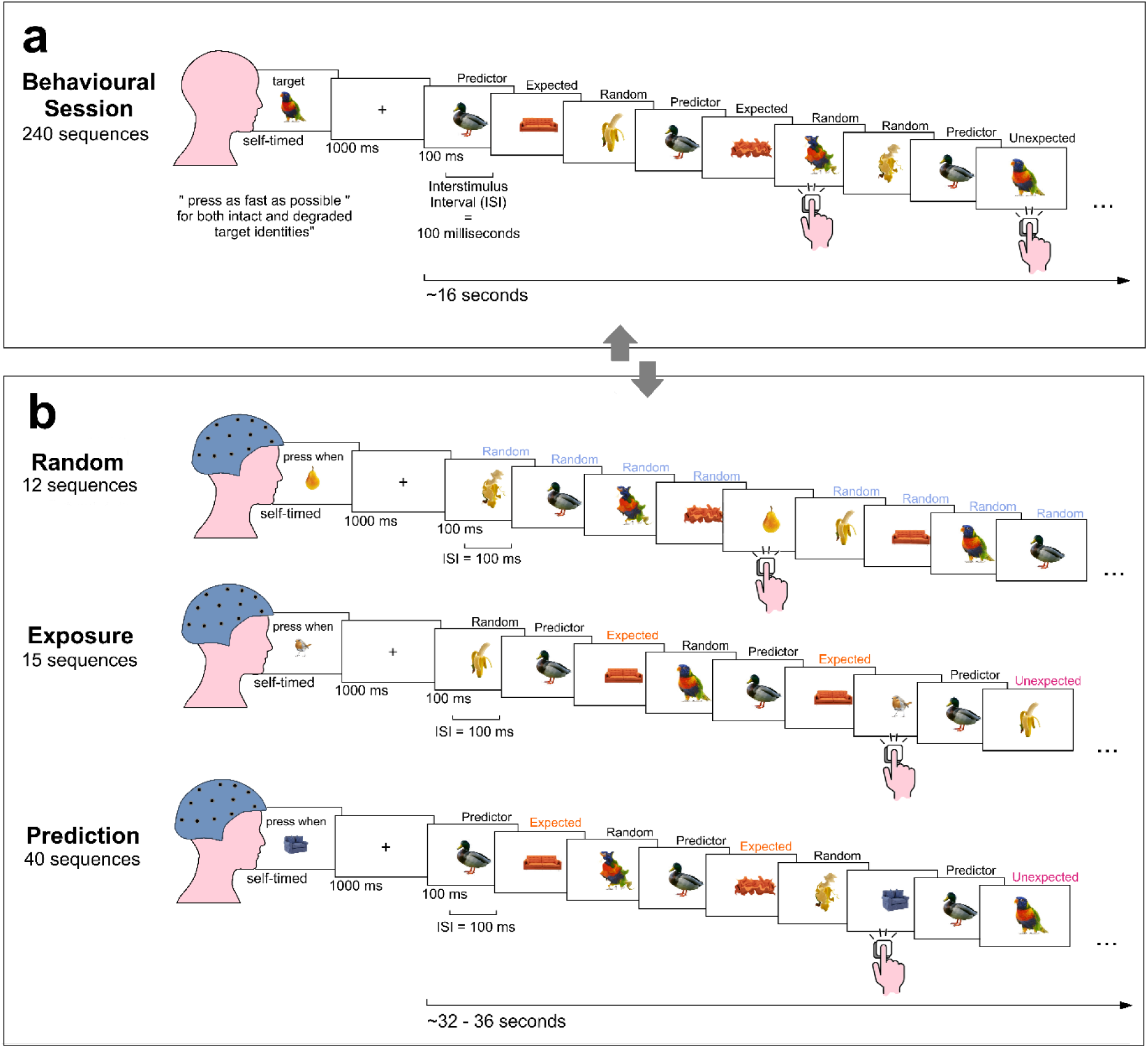
Experimental design with counterbalanced behavioural and EEG sessions. **(a)** The behavioural target detection task. Participants maintained fixation and viewed probabilistically structured sequences, providing speeded responses when a pre-defined target stimulus (degraded and intact) was displayed. **(b)** EEG Random Block sequences contained intact and degraded stimuli presented in random order. **(c)** EEG Exposure Block sequences contained probabilistically structured, intact stimuli. **(d)** EEG probabilistically structured Prediction Block sequences. All sequences included attention probes (one unique identity per sequence).

At the start of the session, participants were shown all intact stimuli and an example of corresponding degraded stimuli. Before each sequence, participants saw an intact target object and were asked to provide a keypress response as quickly as possible when the target (whether intact or degraded) was displayed. Targets were always stimuli which occupied expected, unexpected, and/or random sequence positions. There were 8 targets per sequence (intact and degraded inclusive). Participants completed a practice behavioural task with 6 probabilistically structured sequences using a different stimulus set (Fig.S2). Participants repeated the practice task until they could reliably identify targets (proportion correct ≥ 80%; *M* = 1.05 repeats, *SD* = 0.22).

Responses were considered correct if provided > 150 ms after target presentation, but prior to the interstimulus blank before the next target which occurred at least 500 ms later. This timeframe corresponds with the typical time course for bottom-up visual processing and stimulus-driven behavioural responses (Carlson et al., 2013; Thorpe et al., 1996). We further filtered correct responses to retain only those within 2 *SD* from participants’ overall average reaction time. Target misses occurred when no response was provided within the designated post-target timeframe. False alarms occurred when responses were provided outside the post-target timeframe (i.e., in response to non-target stimuli). Reaction times were measured as the interval between target onset and a correct response as defined above.

### EEG Paradigm

The EEG paradigm evaluated whether expectancy modulates stimulus decoding performance. There were three sequential blocks of sequences (Fig.2). First, the *Random Block* was used to train machine learning classifiers in discriminating stimuli using the associated patterns of neural activity, but in the absence of probabilistic information. Each stimulus (both intact and degraded) was presented 20 times in random order without immediate repeats (12 sequences, > 200 total displays per stimulus). Then, the *Exposure Block* was used to expose participants to stimuli presented within the probabilistic structure before measuring how expectancy modulated neural response outcomes. This block used only intact stimuli (15 sequences, > 300 total displays per predictive pair). Data from this block were not used in subsequent analyses. Finally, the *Prediction Block* was used to determine whether expectancy modulates stimulus decoding accuracy for both intact and degraded stimuli. This block was identical to the Exposure Block except that expected and unexpected stimuli were degraded in 50% of presentations (40 sequences, > 800 total displays per predictive pair).

To maintain participants’ attention during the EEG session, we used the same target detection task employed in the behavioural paradigm, except that the target identities were different. Specifically, EEG sequences contained 3–4 randomly interspersed targets (one unique object per sequence) that were not part of any predictive stimulus set (i.e., ‘attention probes’; Fig.1d). The session differences were adopted to maximise power for detecting behavioural effects (Bays et al., 2015), and to maximise the number of EEG trials which were not affected by motor-related responses (e.g., muscle artefacts and motor preparation activity). Muscle artefacts can mask neural response differences between expectation conditions, while enhanced motor preparation corresponding with a specific stimulus condition can be exploited during decoding to inflate that condition’s accuracy (Grootswagers et al., 2017).

### Debriefing

Debriefing questions were administered after the final testing session to assess participants’ awareness of predictive relationships among the stimuli. First, participants were presented with all intact stimuli. They were asked to connect ‘predictor’ with ‘expected’ stimuli (free response). Next, each predictor stimulus was presented, and participants were asked to choose the stimulus it predicted from a list (multiple choice).

### EEG Decoding Scheme

We used pairwise decoding accuracy over time to measure stimulus-specific neural representations under different expectancy conditions (random, expected, unexpected) and image quality (intact, degraded). Decoding was applied to whole-brain EEG data using CosMoMVPA functions to generate timeseries decoding accuracy (Oosterhof et al., 2016). Specifically, for every participant, regularised linear discriminant analysis classifiers were trained and tested to discriminate neural data associated with each possible stimulus pair per expectancy condition. This occurred for each timepoint independently within the stimulus epoch. Model training and testing was implemented using a pairwise classification scheme (chance accuracy = 50%) and 5-fold cross validation (80/20 split; Grootswagers et al., 2017). The average accuracy across all possible stimulus pairs comprised the decoding performance per expectancy condition, per participant. All classifiers were trained using intact stimulus epochs from the Random Block, which contained no probabilistic structure. To identify differences across expectancy conditions and image quality, these classifiers were tested on data from unexpected and expected stimulus epochs from the Prediction Block, separately for intact and degraded stimuli. Statistical analyses were conducted for each condition using average decoding performance at the group-level.

### EEG Acquisition

We followed the neural acquisition procedure outlined in Moore et al. (2024) Continuous brain activity was recorded using a BioSemi 64-channel system with a 1024 Hz sampling rate, driven right leg and common mode sense electrodes, referenced offline to both mastoids. Electrode arrangement followed the international 10–10 standard (Oostenveld & Praamstra, 2001). EEG data were preprocessed in MATLAB using EEGLAB functions (Delorme & Makeig, 2004). Specifically, we applied high pass (0.1 Hz) and low pass (100 Hz) frequency filters to the raw EEG data. We next identified and rejected noisy electrode channels using joint probability (i.e., channel activations > 5 *SD* from mean) and visual inspection of evoked response potentials (ERPs). Rejected channels were reconstructed via spherical interpolation (*M* = 1.98 channels interpolated, *SD* = 1.68, range = 0–7). In contrast to Moore et al. (2024), we conducted automated independent component analysis (ICA) using the default Luca Pion-Tonachini plugin (Pion-Tonachini et al., 2019), removing components flagged as at least 80% likely to be muscle or eye-movement artifacts (*M* = 4.30 components removed, *SD* = 2.34, range = 1–12). Supplementary analyses indicated that the addition of artefact rejection did not change the qualitative pattern of decoding results (Fig.S3). Data were then down-sampled to 256 Hz, divided into epochs relative to stimulus presentation [-100 ms to +1000 ms], linearly detrended, and baseline corrected.

### Statistical Inference

We computed Bayes Factors (BF_10_) using a Jeffreys-Zellner-Siow (JZS) prior with a default Cauchy distribution (scaling factor = 0.707) to determine evidence for the alternative hypothesis (Jeffreys, 1961; Rouder et al., 2009; Zellner & Siow, 1980). BF_10_ > 3 and BF_10_ > 10 were considered as moderate and strong evidence for the alternative hypothesis, respectively (Morey & Rouder, 2011).

### Data Availability

All experimental code, materials, and processed source data associated with this project are available at doi.org/10.17605/OSF.IO/37FYQ.The experimental design and main analyses were pre-registered prior to data analysis at doi.org/10.17605/OSF.IO/4UQCK. Additional exploratory analyses, which were not pre-registered, are identified as such in the Results section.

## Results

### Expected Stimuli Yielded Faster and Greater Proportion Correct in the Behavioural Task

The proportions of correct responses on the behavioural task were modulated by expectancy (Fig.3a). Expected stimuli yielded the highest proportion of correct responses relative to random and unexpected stimuli. Random and unexpected stimuli did not differ from one another. This was confirmed with a repeated-measures BF Linear Model with participant as a random factor and an BF Inclusion Analysis to quantify the contribution of each model parameter (Clyde et al., 2011; Hinne et al., 2020; Morey & Rouder, 2011). Expectancy yielded moderate evidence of being included in the model (BF_10_ = 7.85). Follow-up pairwise Bayesian *t*-tests (Rouder et al., 2009) confirmed moderate evidence that expected stimuli (*M* = 0.908, *SD* = 0.069) yielded a higher proportion of correct responses relative to random (BF_10_ = 5.38; *M* = 0.894, *SD* = 0.067, *d* = 0.32) and unexpected stimuli (BF_10_ = 5.39; *M* = 0.890, *SD* = 0.076, *d* = 0.32). Random and unexpected stimuli showed moderate evidence against unequal proportions of correct responses (BF_10_ = 0.25).

**Figure 3:**
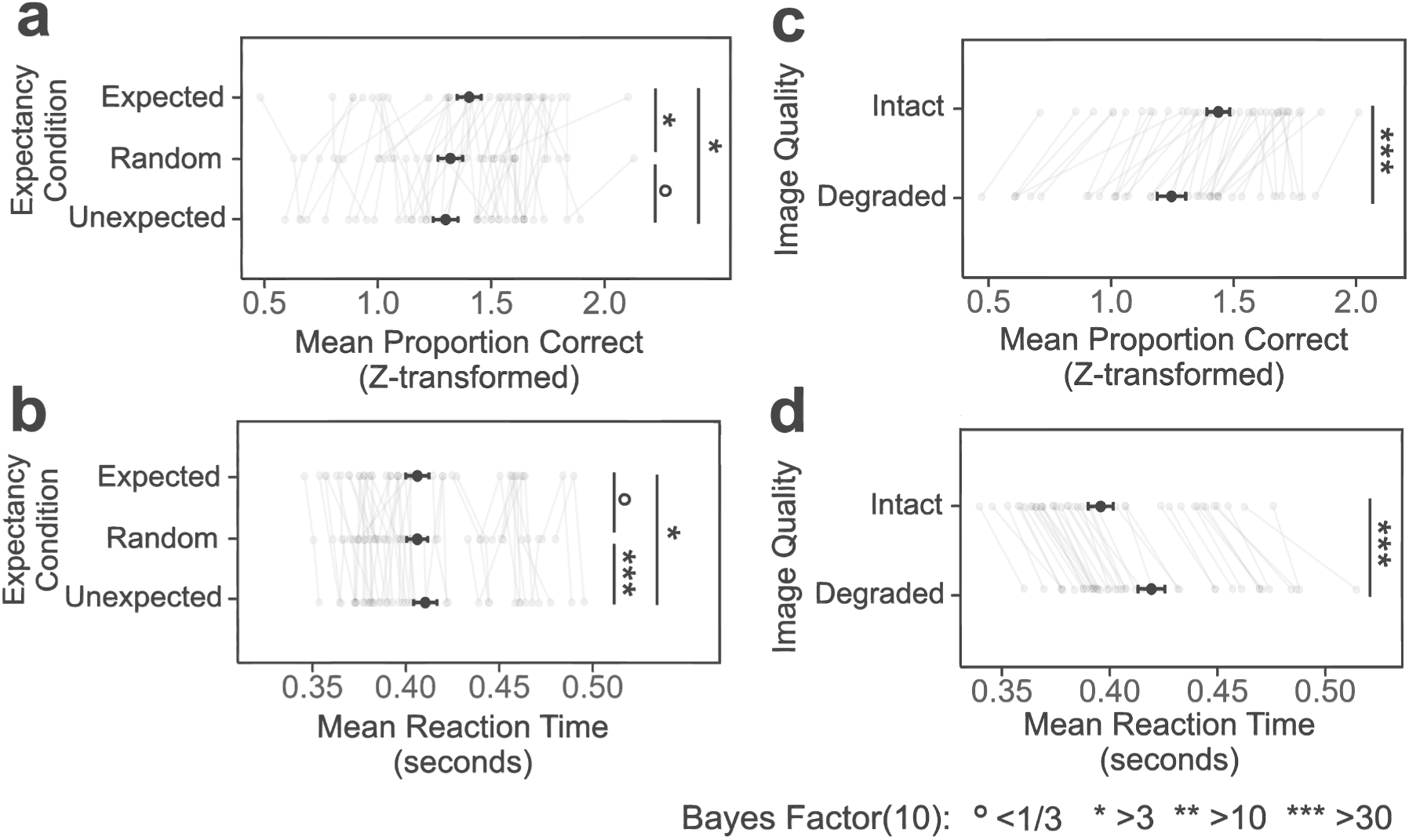
Reaction time and accuracy results from the behavioural target-detection task. **(a)** Proportion of correct responses (Z-transformed to illustrate differences near ceiling) as a function of expectancy and image quality conditions. Statistical tests were conducted using the raw proportion correct. **(b)** Reaction time for correct responses in each expectancy and image quality conditions. Grey lines visualise within-participant condition differences (N = 40) and error bars report standard error of the mean (SEM). BF_10_ ◦ < 1/3, * > 3, ** > 10, *** > 30. Expected and Unexpected Stimuli Yielded Reduced Neural Decoding Accuracy:

Reaction times based on correct responses were also modulated by expectancy. Expected stimuli and random stimuli did not differ, but unexpected stimuli yielded slower reaction times relative to both random and expected stimuli (Fig.3b). Here, expectancy yielded strong evidence of being included in the model (BF_10_ = 11.78). Follow-up comparisons confirmed moderate to strong evidence that unexpected stimuli (*M* = 0.410 s, *SD* = 0.041) yielded slower reaction times relative to both random (BF_10_ = 538.37; *M* = 0.406 s, *SD* = 0.038, *d* = 0.49) and expected stimuli (BF_10_ = 4.46; M = 0.406 s, SD = 0.041, d = 0.31). Random and expected stimuli showed moderate evidence against unequal reaction times (BF_10_ = 0.12).

Notably, expectancy modulations of behavioural measures emerged without participants being able to explicitly report probabilistic transitions among stimuli. There was no evidence that performance on the free response and multiple-choice debrief questions differed from chance (free response: χ^2^(1) = 2.09, BF_10_ = 2.04; multiple-choice: χ^2^(1) = 1.78, BF_10_ = 2.14; Tab.S1,S2).

Finally, relative to intact stimuli, degraded stimuli yielded lower proportion of correct responses (BF_10_ = 7.61e+47) and slower reaction time (BF_10_ = 7.61e+47; Fig.3c,d). Prediction effects did not vary across image quality. There was strong (BF_10_ = 0.08) and moderate (BF_10_ = 0.11) evidence against an expectancy x image quality interaction for reaction times and proportion correct, respectively.

Participants performed the EEG task reliably, providing correct responses to 90.5% of attention-probe targets on average (*SD* = 0.067, range = 68.9%–96.3%). To investigate how expectancy modulates stimulus representation, we examined decoding differences for stimuli when they were random versus when they were expected or unexpected. Classifiers trained to discriminate random stimuli presented in the Random Block were cross-validated and tested on the same stimulus identity (e.g., cow vs. cow) occurring at expected or unexpected sequence positions in the Prediction Block. All model training and testing data can be found in Tab.S3. Here, we report results for intact and degraded stimuli at the group-level (*N* = 40; Fig.4). Overall, regardless of image quality, there was consistent evidence that decoding accuracy for both expected and unexpected stimuli was reduced relative to decoding accuracy for random stimuli.

**Figure 4:**
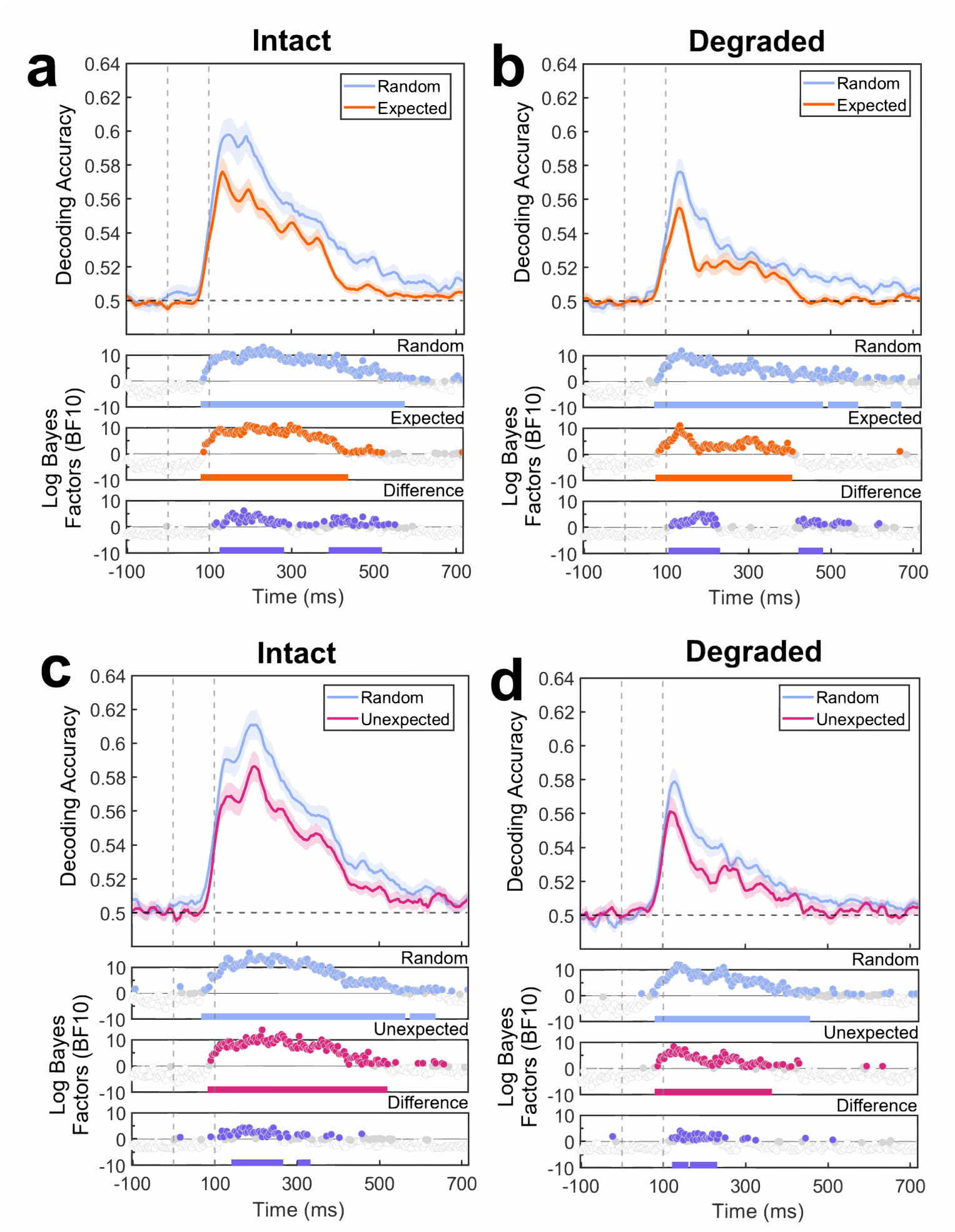
Decoding results for expected and unexpected stimuli across image quality conditions. **(a) Top panel:** Decoding accuracy for intact random stimuli (Random Block) and expected stimuli (Prediction Block). Stimulus onset was at 0 milliseconds (ms) and offset was at 100 ms. Shaded regions are standard error of the mean (SEM). **Bottom panels:** Bayes Factors (BF_10_) show evidence for above-chance decoding and condition differences (purple panel). BFs y-axes are exponents of 10. Solid coloured lines underneath the BF timeseries highlight timepoints surviving cluster-based permutation tests (p < .05). **(b)** Decoding accuracy and BF timeseries for degraded random stimuli (Random Block) and expected stimuli (Prediction Block). **(c)** Decoding accuracy for intact random stimuli (Random Block) and unexpected stimuli (Prediction Block). **(d)** Decoding accuracy for degraded random stimuli and unexpected stimuli.

Evidence for or against above-chance decoding accuracy and between-model differences were evaluated at each timepoint using Bayesian *t*-tests. We employed a null interval of 0–0.5 effect size to account for potential classifier biases, one-sample tests for comparing to chance, and two-sample tests for comparing between conditions. Above-chance decoding onset was designated as the first of at least three consecutive timepoints across which there was moderate evidence (BF_10_ > 3) for above-chance decoding (Mai et al., 2019). Timepoints yielding BF_10_ > 3 were also included in clusters. We applied frequentist cluster-based permutation to correct for multiple comparisons by calculating the probability that each observed cluster size occurred by chance. Clusters occurring with *p* < 0.05 after 10,000 permutations were considered significant. All significant timepoint clusters are reported relative to stimulus onset (0 ms). We focus on clusters between 0–600 ms because this is the time window that captures the typical emergence and decay of object-level information (Carlson et al., 2013), and where prediction effects have been found previously (Moore et al., 2024; Tang et al., 2018).

Across all timeseries, above chance decoding first emerged between 74 to 90 ms, which is typical for object-level decoding (Carlson et al., 2013; Grootswagers et al., 2019). However, expected stimuli yielded significantly lower decoding accuracy relative to random stimuli (Fig.4a,b). For intact stimuli, this difference was statistically significant between, 133– 273 ms (37 timepoints), 398–414 ms (5 timepoints), 422–453 ms (9 timepoints), 465–481 ms (5 timepoints) and 496–512 ms (5 timepoints). For degraded stimuli, the same effect was observed between 113–222 ms (29 timepoints), 429–453 (7 timepoints), and 460–472 (4 timepoints). Likewise, unexpected stimuli yielded significantly lower decoding accuracy relative to random stimuli (Fig.4c,d). For intact stimuli, this difference was statistically significant between 148–195 ms (13 timepoints), 203–258 ms (15 timepoints), and 309–324 ms (5 timepoints). For degraded stimuli, the difference was significant between 128–152 ms (7 timepoints) and 171–222 ms (14 timepoints). Notably, the reduced decoding performance for expected and unexpected stimuli across image quality conditions is not explained by the pattern of evoked response potentials (ERPs) as characterised by peak-to-peak amplitude analyses (Fig.S4,S5).

### Prior Exposure Did Not Change the Observed Behavioural and Neural Effects

In our study, participants who completed either the EEG or behavioural task in the second session had prior exposure to stimuli presented in predictive relationships (during the previous testing session). Conversely, participants who completed either the EEG or behavioural task in the first session would not have had such exposure. To determine whether prior exposure influences prediction effects in behavioural performance and neural decoding, we conducted exploratory (un-preregistered) analyses by splitting participants into Prior Exposure and No Prior Exposure groups (*n* = 20 each) and then repeating the behavioural and neural analyses.

Prior exposure did not change the pattern of neural or behavioural effects observed in the group-level analyses. A reanalysis of behavioural measures using BF Linear Models yielded evidence against including prior exposure, whether as a modulator of previously observed behavioural results for expectancy (reaction time: BF_10_ = 0.07; proportion correct: BF_10_ = 0.09), image quality (reaction time: BF_10_ = 0.46; proportion correct: BF_10_ = 0.21), or the expectancy x image quality interaction (reaction time: BF_10_ = 0.11; proportion correct: BF_10_ = 0.25).

For the neural data, decoding accuracy for expected and unexpected stimuli was reduced relative to random stimuli across all image quality conditions and prior exposure groups. Although this trend was the same across exposure groups, however, they were not identical. Relative to the Prior Exposure group, No Prior Exposure participants showed reduced decoding over a small temporal range for expected stimuli (6 timepoints for largest cluster from 80–85 ms, Fig.5a), and a large temporal range for unexpected stimuli (11 timepoints for largest cluster from 65–75 ms, Fig.5c). In contrast, Prior Exposure participants showed reduced decoding over a relatively larger temporal range for expected stimuli (31 timepoints for largest cluster from 62–92 ms, Fig.5b), but over a smaller temporal range for unexpected stimuli (5 timepoints for largest cluster from 86–90 ms, Fig.5d). These effects were consistent across intact and degraded stimuli (Fig.S6), suggesting an important influence of prior exposure on the extent of neural prediction effects.

**Figure 5:**
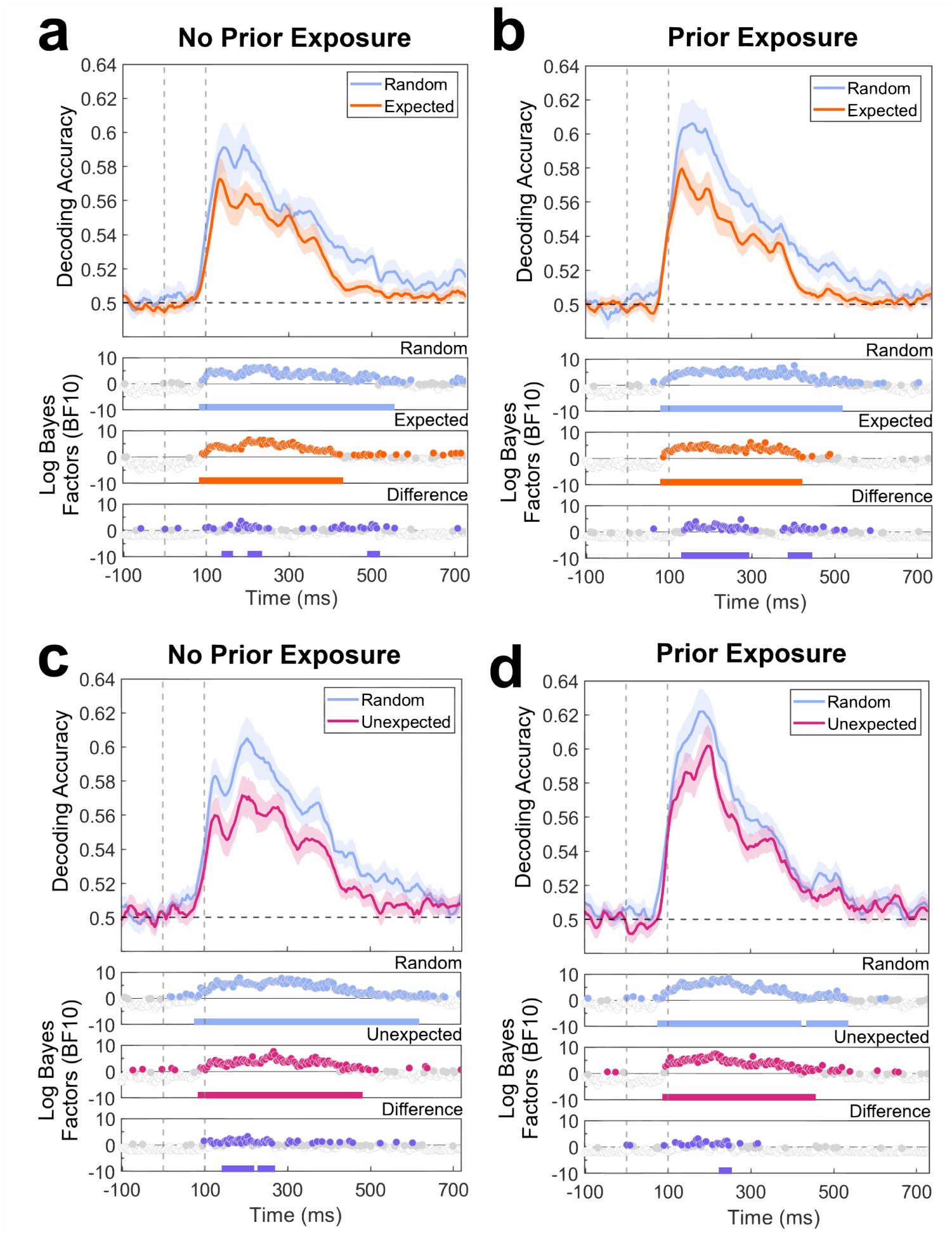
Decoding results for intact expected and unexpected stimuli across prior exposure groups. **(a) Top panel:** Decoding accuracy for intact random stimuli (Random Block) and expected stimuli (Prediction Block) for No Prior Exposure participants (n = 20). Stimulus onset was at 0 milliseconds (ms) and offset was at 100 ms. Shaded regions are standard error of the mean (SEM). **Bottom panels:** Bayes Factors (BF_10_) show evidence for above-chance decoding and condition differences (purple panel). BFs y-axes are exponents of 10. Solid coloured lines underneath the BF timeseries highlight timepoints surviving cluster-based permutation tests (p < .05). **(b)** Decoding accuracy and BF timeseries for intact random stimuli (Random Block) and expected stimuli (Prediction Block) for Prior Exposure participants (n = 20). **(c-d)** Decoding of intact random stimuli (Random Block) and unexpected stimuli (Prediction Block) for No Prior Exposure and Prior Exposure participants. See Supplementary (Fig.S4) for degraded stimulus decoding across prior exposure groups.

### Prior Exposure was Associated with Pre-Activation of Expected Information even in the Absence of Predictive Relationships

We also asked whether prior exposure had other effects on neural activity that was not characterised by the previous decoding analyses. The neural representation of expected visual stimuli and their preceding predictor stimuli have been shown to merge following exposure to probabilistic relationships (Schapiro et al., 2012). This suggests a shared representation between sequential stimuli with repeated pairings, and the possibility that expected stimuli are represented prior to stimulus onset.

To determine whether expected stimuli are represented before stimulus onset and how this may be modulated by prior exposure, we conducted exploratory (un-preregistered) cross-decoding analyses (e.g., Moore et al., 2024). Specifically, we asked whether neural responses following the display of predictor stimuli contained information about the upcoming (but not yet displayed) expected stimulus in Prior Exposure and No Prior Exposure participants.

Classifiers were trained to discriminate both intact and degraded stimuli from the Random Block, which contained no predictive information (Fig.6a). These models were then tested on Prediction Block epochs containing each stimulus’ associated predictor stimuli (hereafter called the ‘Recoded Predictor Model’). Above-chance decoding in this Recoded Predictor Model would suggest that neural activity associated with predictors contains stimulus-related information about the subsequent – but not yet displayed – expected stimuli. To ensure any above-chance decoding was not solely driven by visual or semantic similarities between predictor and expected stimuli, classifiers were also tested on predictors occurring randomly in the Random Block before predictive relationships were established (hereafter the ‘Recoded Control Model’). Given that no predictive relationships were present in these trials, the control model should perform at chance.

**Figure 6:**
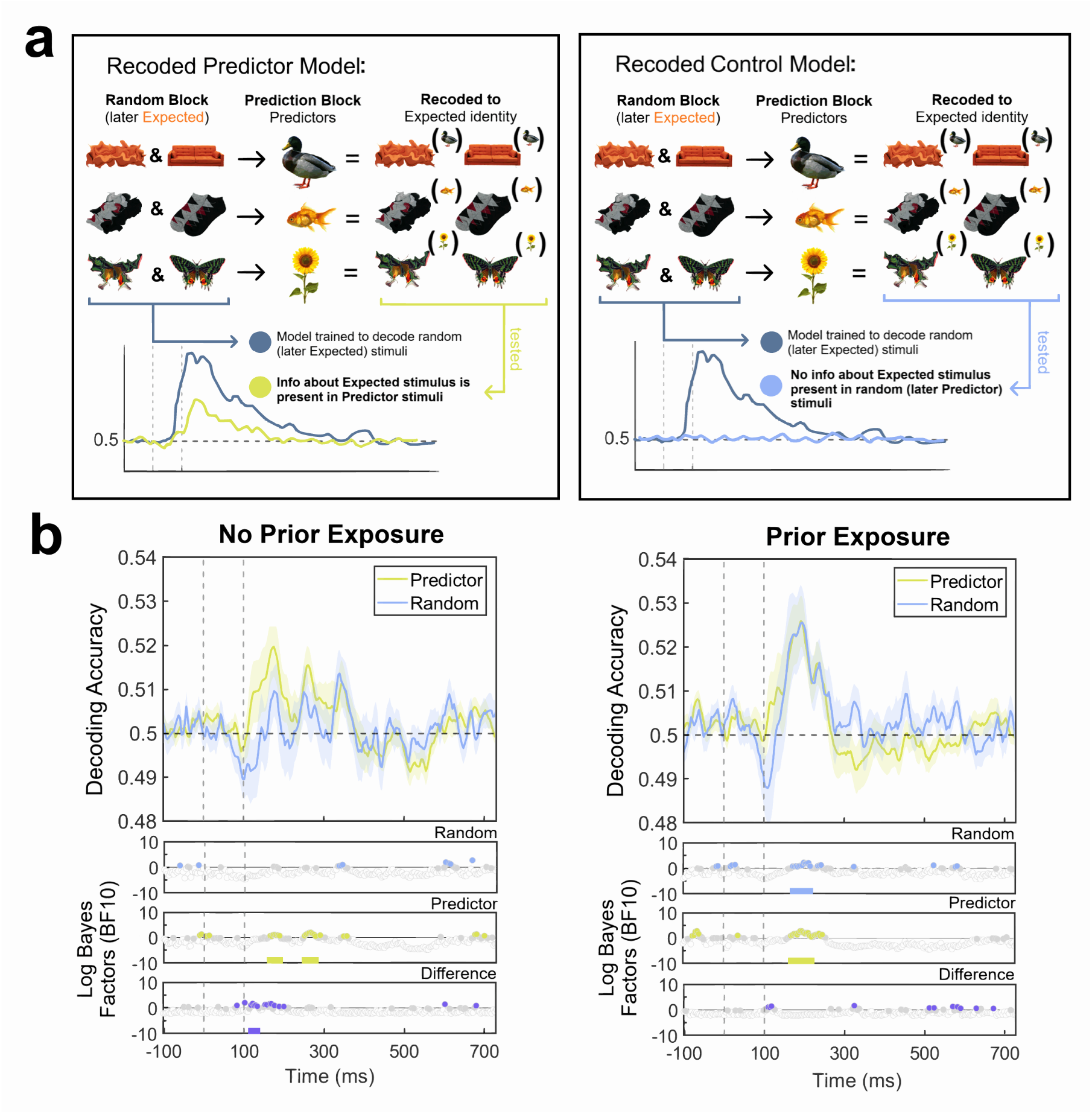
Cross-decoding model construction and results assessing whether predictors were decoded above chance using classifiers trained on Random Block (later expected) stimuli, split by prior exposure. **(a)** In the Recoded Predictor Model, the identity of expected stimuli was decoded from Prediction Block trials in which predictor stimuli were presented. In the Recoded Control Model, the identity of expected stimuli was decoded from Random Block trials in which random (later predictor) stimuli were presented. Both of these models were tested against classifiers trained to decode random (later expected) stimuli from the Random Block. **(b) Top panels:** Decoding accuracy of predictor stimuli in the Recoded Control and Predictor Models for No Prior Exposure participants and Prior Exposure participants (n = 20 each). Stimulus onset was at 0 milliseconds (ms) and offset was at 100 ms. Shaded regions are standard error of the mean (SEM). **Bottom panels:** Bayes Factors (BF_10_) show evidence for above-chance decoding and condition differences (purple panel). BFs y-axes are exponents of 10. Solid coloured lines under the BF timeseries highlight timepoints surviving cluster-based permutation tests (p < .05).

The results suggest that predictors contained stimulus-relevant information about the subsequent expected stimuli (Fig.6b). For No Prior Exposure participants, decoding accuracy for predictor stimuli was at chance level for the Control Model. By contrast, for the Recoded Predictor Model, decoding accuracy was above chance at 164–188 ms (7 timepoints) and 250–277 ms (8 timepoints). For Prior Exposure participants, both the Control Model and Recoded Predictor Model yielded above chance decoding accuracy, respectively from 172–214 ms (12 timepoints) and 168–219 ms (14 timepoints). It is notable that the Control Model performed above chance here, indicating that prior exposure to predictive relationships may influence neural responses in the Random Block. Taken together, regardless of prior exposure, information about subsequent expected stimulus is decodable during the time interval of the preceding predictor stimulus. With prior exposure, this information is also available even when no predictive structure underlies stimulus presentations.

## Discussion

This study investigated how prior exposure to probabilistic relationships and image quality modulate prediction effects in neural and behavioural measures of object recognition. In the detection task, participants showed faster response times and higher accuracy for expected stimuli relative to unexpected stimuli. However, this effect was not modulated by image quality or prior exposure. Both expected and unexpected stimuli yielded reduced decoding accuracy relative to corresponding random stimuli. This pattern was consistent across image quality conditions. Exploratory analyses nevertheless provided suggestive evidence that subtle representational changes may be linked to prior exposure. Taken together, the results suggest that prior exposure and image quality do not play influential roles in modulating prediction effects during visual object recognition.

### Prediction Effects in Neural and Behavioural Measures of Object Processing

Expected stimuli yielded a higher proportion of correct responses relative to unexpected and random stimuli, but there was no difference between unexpected and random stimuli. Expected stimuli also yielded faster reaction times than unexpected stimuli. These results align with previous studies reporting that expected objects placed within probabilistic pairs or sequences are detected faster and more accurately relative to random and/or unexpected stimuli (Pinto et al., 2015; Stein & Peelen, 2015; Turk-Browne et al., 2005). The results of the current study align with this past work in suggesting that expected stimuli confer a detection performance advantage.

Multivariate analyses of EEG activity showed that both expected and unexpected stimuli yielded reduced decoding performance relative to random stimuli. This finding aligns with past studies which also reported reduced neural response amplitudes and representational fidelity using probabilistic image sequences (Kumar et al., 2017; Richter et al., 2018, 2024). More broadly, the findings align with a hallmark of predictive processing known as expectation suppression, which is assumed to occur when an expectation is fulfilled (de Lange et al., 2018; Summerfield et al., 2008). However, our results suggest that this dynamic may not be exclusive to expected stimuli because unexpected stimuli also yielded lower decoding accuracy relative to random stimuli. Other EEG studies employing probabilistic image sequences have also reported evidence for equal, or comparatively reduced decoding accuracy, in unexpected relative to expected and random object stimuli (den Ouden et al., 2023, 2025; Moore et al., 2024). Our results agree with past work indicating that the occurrence and directionality of neural prediction effects may vary across different experimental paradigms.

### Image Quality Does Not Modulate Behavioural and Neural Prediction Effects

In the current study, prediction effects measured in behavioural and neural responses were consistent across image quality conditions. The observed advantage in detecting expected stimuli aligns with past contextual prediction research reporting that objects in semantically congruent scenes are detected more accurately regardless of image quality (Brandman & Peelen, 2017). Expectation fulfillment thus benefits object detection performance even when not all object-level information is available due to degraded image properties. It is worth noting that images in the current study were degraded to a level intended to reduce, but not eliminate, object recognisability (Stojanoski & Cusack, 2014). It is possible that the magnitude of detection performance benefits would be reduced with more dramatic image degradation. The current behavioural findings nevertheless imply that greater object-specific information may be present in neural activity when expectations are fulfilled relative to when there is no expectation (i.e., random images), as suggested by Moore et al. (2024). In contrast, we found that expected stimuli were decoded more poorly relative to random stimuli, and this difference did not change with image quality. Overall, our results suggest that the current image quality manipulation did not significantly modulate behavioural and neural prediction effects, and that improved behavioural performance does not always correspond with improved fidelity of neural representations.

### Prior Exposure Does Not Modulate Behavioural and Neural Prediction Effects

Prediction effects measured in behavioural and neural responses were consistent across exposure conditions. Previous research has shown that participants can learn probabilistic relationships among sequential stimuli quickly (e.g., within 200 repetitions of a predictive sequence), even if implicitly (Bays et al., 2015; Turk-Browne et al., 2005). This suggests that behavioural prediction effects can be detected shortly after exposure to probabilistic inputs. Our results extend this observation by demonstrating that prior exposure over an extended duration (> 2000 repetitions of probabilistic pairs) does not necessarily confer added detection benefits for expected stimuli.

By contrast, the absence of an effect of prior exposure on neural decoding accuracy was surprising. Despite there being reduced decoding accuracy for expected and unexpected stimuli across prior exposure conditions, exploratory analyses provided tentative evidence that prior exposure may modulate some aspects of neural responses to predictive relationships. Specifically, prior exposure was associated with a reduction in decoding performance over a larger temporal range for expected stimuli, and over a smaller temporal range for unexpected stimuli. This suggests that increased exposure strengthened how long decoding suppression occurred for expected (but not unexpected) stimuli. However, these effects were subtle and insufficient in magnitude to change the group-level effect where expected and unexpected stimuli yielded reduced decoding performance relative to random. We acknowledge that such reduced decoding performance could be due to classifier models being optimised to decode random stimuli. This is unlikely to be the only explanation for the findings, however, because the analysis approach we used is commonly employed for investigating effects of prediction on visual processing (Hogendoorn & Burkitt, 2018; Moore et al., 2024; Richter et al., 2024). In some cases, past work has also found improved decoding of expected stimuli relative to the random baseline (e.g., Moore et al., 2024).

Alternatively, it is possible that the lack of influence by prior exposure is because effects may not be captured by decoding performance after stimulus onset. Exploratory analyses indicated that information about expected stimuli was already represented in neural activity when predictor stimuli were displayed. These results suggest that prediction influences neural responses outside the standard post-onset window of stimulus presentation, in line with several previous studies (Ekman et al., 2017; Kok et al., 2017). Notably, these effects occurred even when there were no predictable relationships (i.e., for random predictor stimuli). Pre-activation did not occur for all random predictor stimuli, but only for those participants who benefitted from prior exposure. In participants without any prior exposure to predictive relationships, there was no expected information present in the neural activity of random predictor stimuli. Thus, the results could not be due to classifier contamination or featural similarity between predictor and expected stimuli alone. Prior exposure to predictive relationships led to the pre-activation of expected stimulus features even within random image sequences in the second session.

It remains unclear, however, which visual or semantic features were responsible for the observed pre-activation effects and whether such features are task specific. Future research should delineate features contributing to above-chance decoding of expected information before stimulus onset by training classification models to parse neural activity based on feature dimensions of increasing complexity (e.g., orientation, colour, contours, category, and animacy; Groen et al., 2017; Grootswagers et al., 2019). Future research should also investigate whether the contributing features are specific to task requirements; a colour classification versus animacy classification task may demand that participants prioritise different features of the same image.

Moreover, the occurrence of predictive stimulus pre-activation may itself be task-dependent and related to the superior detection of expected stimuli. If pre-stimulus information is task-relevant, response preparation may occur earlier in time (Mulder et al., 2012; Thakur et al., 2021; Turk-Browne et al., 2010), leading to rapid and more accurate behavioural performance when confirmed by the expected stimulus presentation. Future research should evaluate how predictive signals about upcoming expected stimuli are related to behavioural performance, and whether this is modulated by prior exposure and image quality.

## Conclusion

Here, we asked whether image quality and prior exposure impact neural and behavioural prediction effects. Although we observed clear evidence of predictive effects on behaviour, the neural decoding analyses provided no clear evidence in line with predictive processing. Critically, image quality and prior exposure did not modify the observed behavioural and neural prediction effects, though there was preliminary evidence that prior exposure may subtly change the representational time course of expected stimuli. Our study extends current understanding about the role of prediction during object recognition and contributes to the continuing debate regarding how the brain utilises predictive relationships to support perception.

## CReDiT statement

**Phuong N.U. Dang:** Conceptualisation (30%), Methodology (30%), Software (95%), Data curation, Formal analysis, Investigation (50%), Visualisation, Writing – Original Draft. **Jason B. Mattingley:** Conceptualisation (35%), Methodology (35%), Resources, Writing – Review & Editing, Funding acquisition. **Margaret J. Moore:** Conceptualisation (35%), Methodology (35%), Software (5%), Writing – Review & Editing, Supervision, Project administration, Funding acquisition.

## Acknowledgement

The authors thank Mr Douglas B. Cribb for his assistance with data collection.

