## Supplementary for "Exploring the role of prior exposure and image quality in neural and behavioural prediction effects"

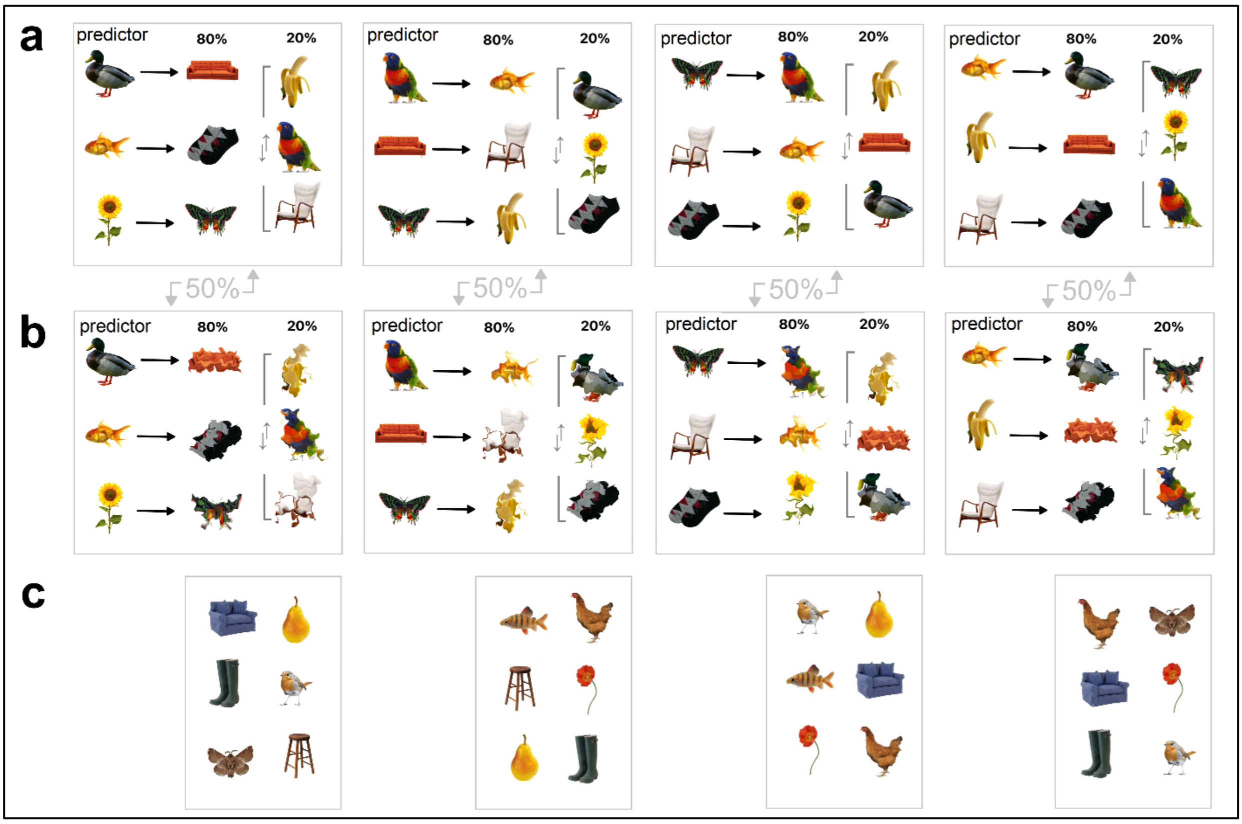


Figure Supplement 1: *All four stimulus sets, one to which a given participant was randomly assigned.* ***(a)*** *Different objects were predictors, expected (80% predicted), and unexpected in each set.* ***(b)*** *The same stimulus set but with degraded objects (one version for EEG, three versions for Behaviour).* ***(c)*** *Attention-probe stimuli for the EEG version of the task.*

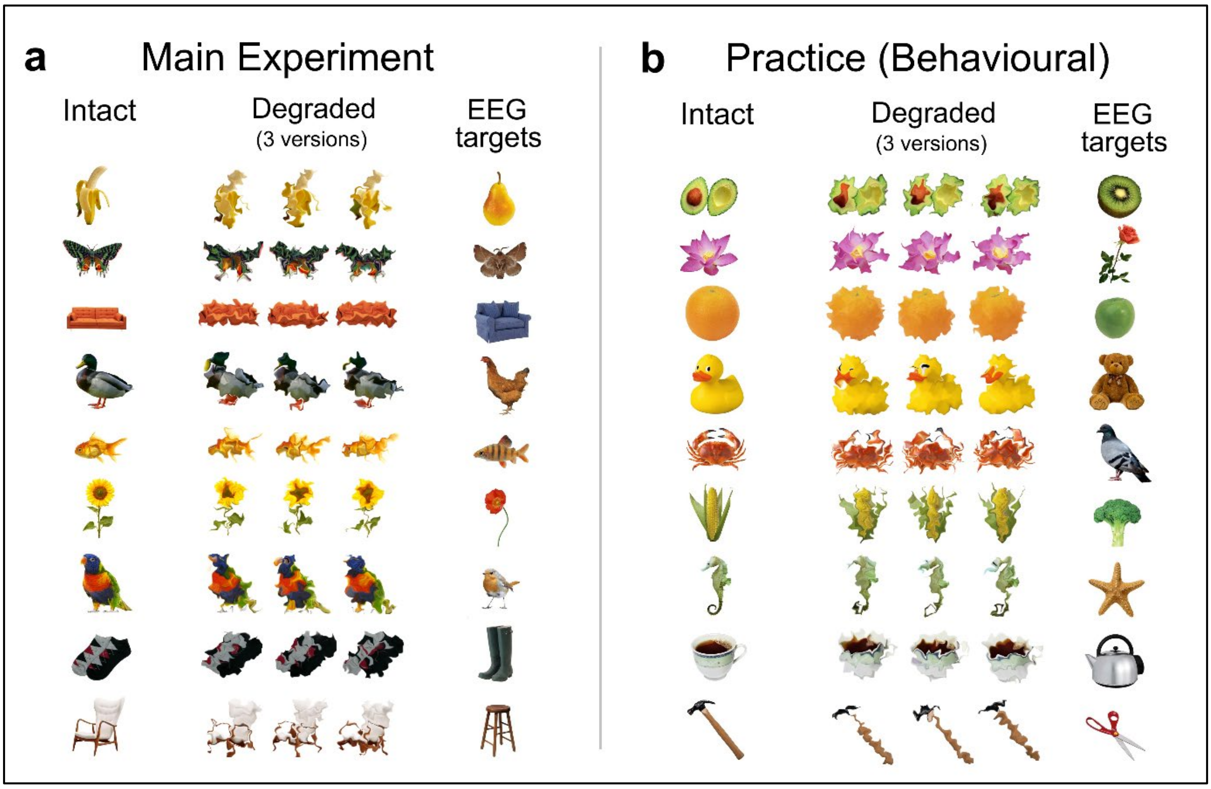


*<u>Figure Supplement 2</u>: All stimuli used in the experiment.* ***(a)*** *Stimuli used in both the neural and behavioural experiments. Degraded stimuli for the behavioural experiment included all 3 versions using different warping matrices implemented at the same level of distortion. Degraded stimuli for the EEG experiment only included the left-most column of warped objects.* ***(b)*** *Stimuli used in the practice version of the behavioural task. Practice stimuli were structured in the same way as the main behavioural experiment.*

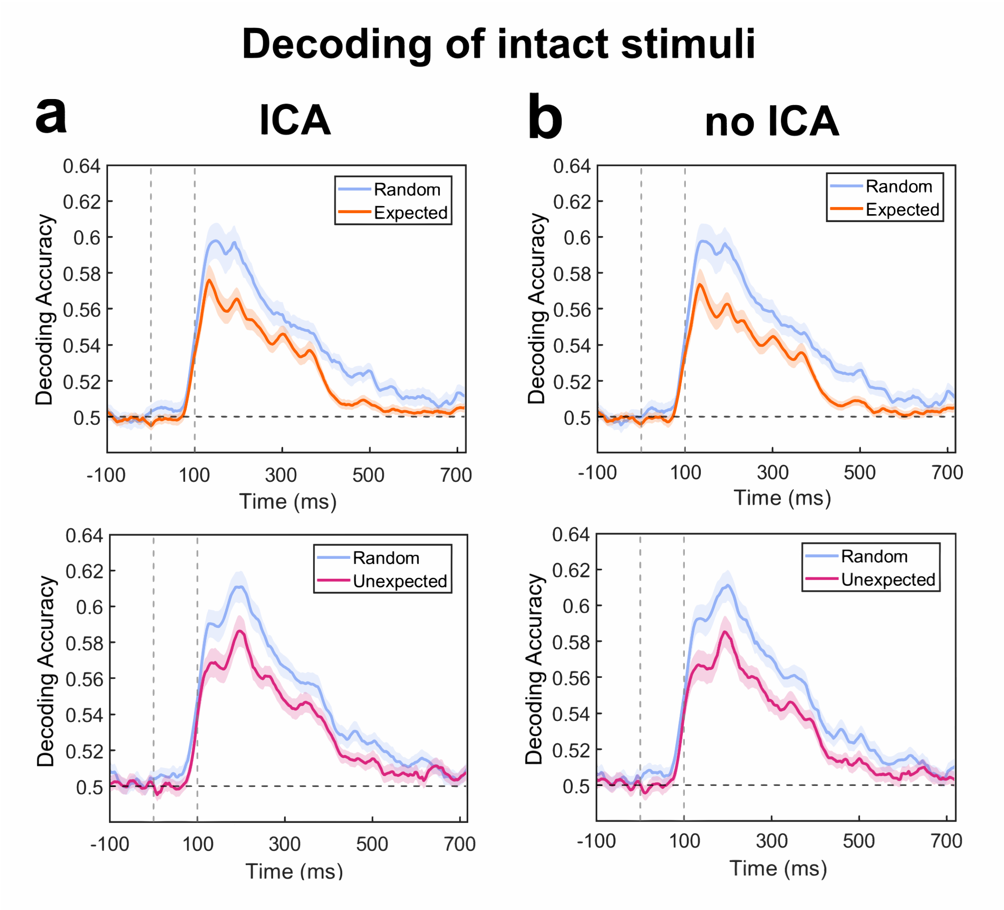


*Figure Supplement 3: Decoding results for intact stimuli with and without ICA.* ***(a)*** *Decoding results for intact random versus expected and unexpected stimuli, as depicted in Figure 3 in the main text. This uses data with ICA to remove components flagged as muscle and eye-movement artefacts.* ***(b)*** *Decoding results for intact random versus expected and unexpected stimuli. This uses data without ICA. Stimulus onset was at 0 milliseconds (ms) and offset was at 100 ms. Shaded regions are standard error of the mean (SEM).*

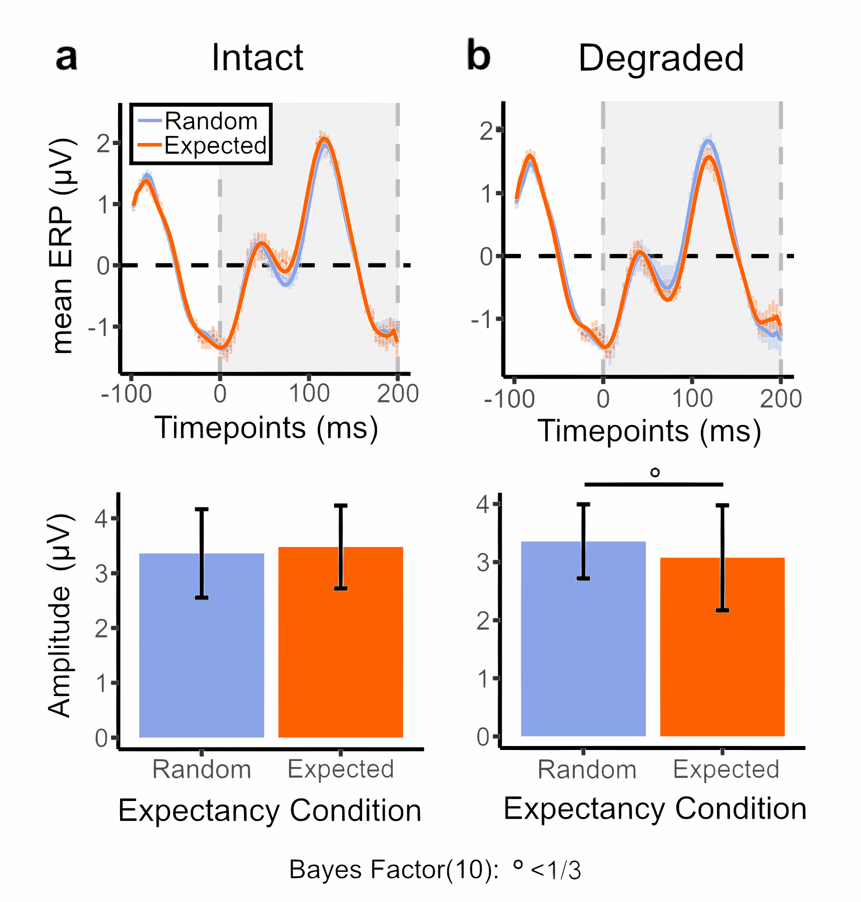


*<u>Figure Supplement 4</u>: Evoked response potential waveforms and peak-to-peak amplitude for random (Random Block) versus expected stimuli (Prediction Block), split by image quality. ERPs are time-locked to stimulus onset at 0 milliseconds (ms). The stimulus onset asynchrony is 200 ms (100 ms stimulus on, 100 ms interstimulus blank).* ***(a) Top:*** *ERP waveform for random and unexpected intact stimuli averaged over the occipital electrode cluster (Oz, O1, O2, POz, PO7, PO3, P5, P7, PO8, PO4, P6, P8; Oostenveld & Praamstra, 2001). Shaded regions are standard error of the mean (SEM).* ***Bottom:*** *Peak-to-peak ERP amplitude for random versus expected stimuli (0-200ms). Error bars report standard error of the mean (SEM).* ***(b) Top and Bottom:*** *ERP waveform and amplitude for degraded stimuli in random versus expected condition.* $BF_{10}$ *◦ < 1/3, unmarked is inconclusive.*

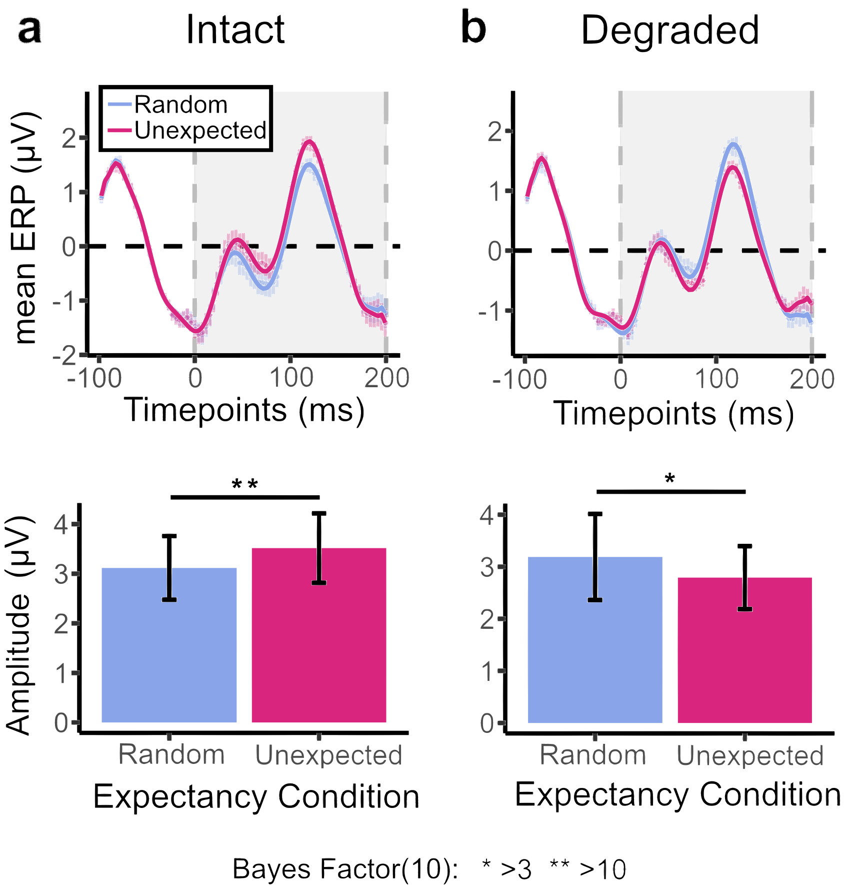


*<u>Figure Supplement 5</u>: Evoked response potential waveforms and peak-to-peak amplitude for random (Random Block) versus unexpected stimuli (Prediction Block), split by image quality. ERPs are time-locked to stimulus onset at 0 milliseconds (ms). The stimulus onset asynchrony is 200 ms (100 ms stimulus on, 100 ms interstimulus blank). **(a) Top:** ERP waveform for random and unexpected intact stimuli averaged over the occipital electrode cluster (Oz, O1, O2, POz, PO7, PO3, P5, P7, PO8, PO4, P6, P8). Shaded regions are standard error of the mean (SEM). **Bottom:** Peak-to-peak ERP amplitude for random versus unexpected stimuli (0-200ms). Error bars report standard error of the mean (SEM). **(b) Top and Bottom:** ERP waveform and amplitude for degraded stimuli in random versus unexpected condition.* $BF_{10}$ ** > 3, ** > 10.*

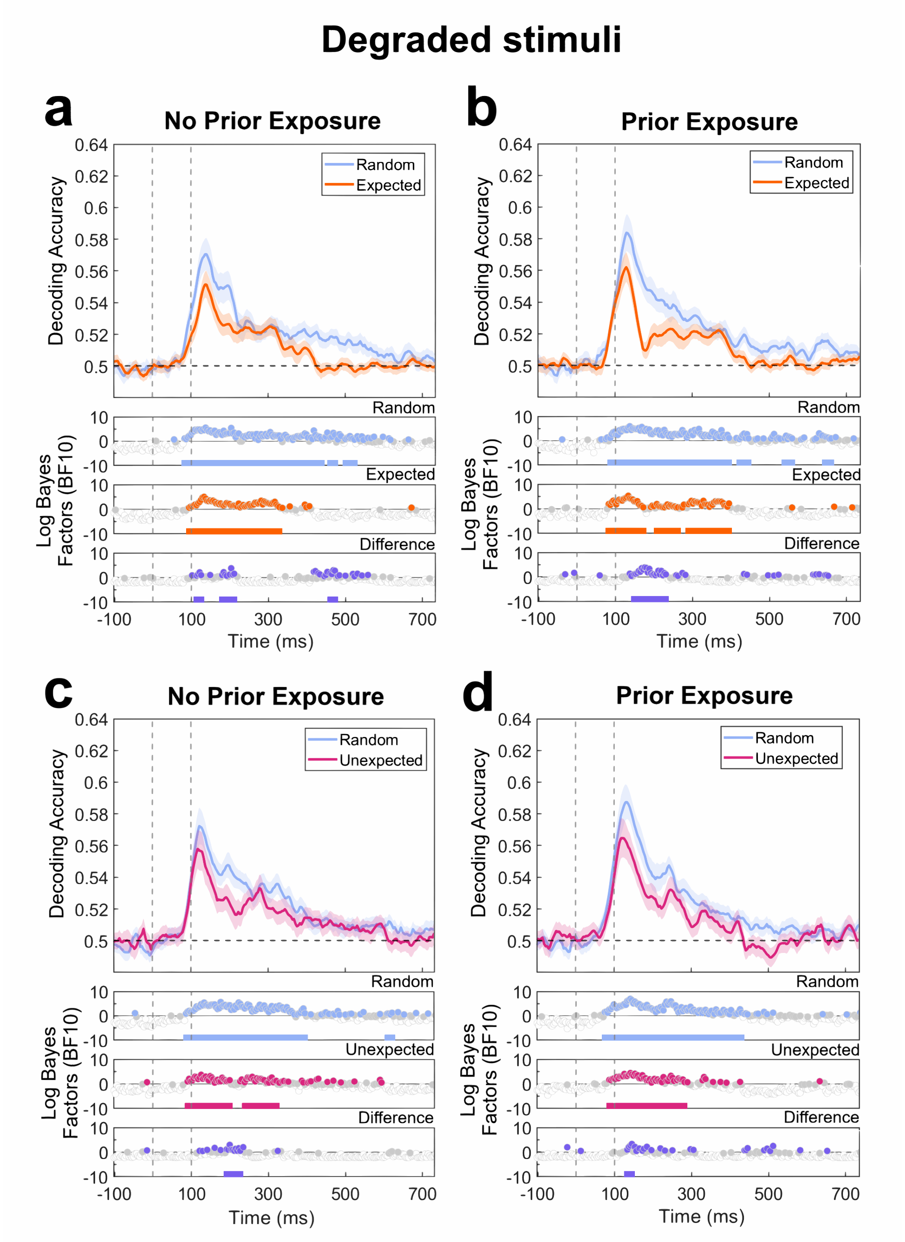


Figure Supplement 6: *Decoding results for degraded expected and unexpected stimuli across prior exposure groups.* ***(a) Top panel:*** *Decoding accuracy of degraded random stimuli (Random Block) and expected stimuli (Prediction Block) for No Prior Exposure participants (n = 20). Stimulus onset was at 0 milliseconds (ms) and offset was at 100 ms. Shaded regions are SEM.* ***Bottom panels:*** *Bayes Factors (BF10) show evidence for above-chance decoding and condition differences (purple panel). BFs y-axes are exponents of 10. Solid coloured lines underneath the BF timeseries highlight timepoints surviving cluster-based permutation tests (p < .05).* ***(b)*** *Decoding accuracy and BF timeseries of degraded random stimuli (Random Block) and expected stimuli (Prediction Block) for Prior Exposure participants (n = 20).* ***(c-d)*** *Decoding of degraded random stimuli (Random Block) and unexpected stimuli (Prediction Block) for No Prior Exposure and Prior Exposure participants.*

| **FreeCh** | expected | observed (average) |
|---|---|---|
| correct pairs | 3 | 0.375 |
| remainder | 78 | 80.625 |
| total possible pairs | 81 (9*9) | 81 (9*9) |

```
Pearson's Chi-squared test with simulated p-value
(based on 1000 replicates)
(test to reject null of non-indep.)
X-squared = 2.0851, df = NA, p-value = 0.2478
--------------
Bayes factor analysis (evidence for non-indep.)
[1] Non-indep. (a=1) : 2.037975 ±0%
Against denominator:
  Null, independence, a = 1
Bayes factor type: BFcontingencyTable,
independent multinomial
```

*<u>Table Supplement 1:</u> Statistical test of independence from chance for the average number of correct responses (across participants, N = 40) on free-choice debriefing questions. The contingency table calculates the proportions of responses given random guessing and the average number of correct responses provided across participants. Note that there are 3 correct pairs of object stimuli with reliable predictive relations. Statistical tests involved both frequentist Chi-square and Bayesian contingency tests.*

| **MCQ** | expected | observed (average) |
|---|---|---|
| correct responses | 3 | 0.575 |
| remainder | 21 | 23.425 |
| total possible responses | 24 ([9-1]*3) | 24 ([9-1]*3) |

```
Pearson's Chi-squared test with simulated p-value
(based on 1000 replicates)
(test to reject null of non-indep.)
X-squared = 1.7773, df = NA, p-value = 0.2378
--------------
Bayes factor analysis (evidence for non-indep.)
[1] Non-indep. (a=1) : 2.136364 ±0%
Against denominator:
  Null, independence, a = 1
Bayes factor type: BFcontingencyTable,
independent multinomial
```

*<u>Table Supplement 2:</u> Statistical test of independence from chance for the average number of correct responses (across participants, N = 40) on the multiple-choice debriefing questions. The contingency table calculates the proportions of responses given random guessing and the average number of correct responses provided across participants. Note that there are 3 correct pairs of object stimuli with reliable predictive relations. Statistical tests involved both frequentist Chi-square and Bayesian contingency tests.*

| Training data | Testing data | Average epoch numbers (SD) | Onset (ms) | Peak (ms) | Peak (%) | BF>3 timepoints in ms [count] | |
|---|---|---|---|---|---|---|---|
| | | | | | | **Above 50%** | **Difference** |
| **Intact stimuli** | **Intact stimuli** | | | | | | |
| | Expected | 177.09 (2.19) | 85.9 | 183.6 | 60.23 | 85–511 | 132–273 [37] |
| Expected | Random Block | 44.12 (0.50) | | | | 519–566 | 398–414 [5] |
| Random Block | | | | | | | 421–453 [9] |
| | expected | 177.09 (2.19) | 74.2 | 191.4 | 61.35 | 85–429 | 464–480 [5] |
| | Prediction Block | 122.03 (0.92) | | | | | 496–511 [5] |
| **Intact stimuli** | **Intact stimuli** | | | | | | |
| | Unexpected | 176.93 (2.73) | 85.9 | 132.8 | 57.80 | 74–554 | |
| Unexpected | Random Block | 44.08 (0.63) | | | | 582–605 | 148–195[13] |
| Random Block | | | | | | | 203–257[15] |
| | Unexpected | 176.93 (2.73) | 89.8 | 199.2 | 59.15 | 89–480 | 308–324 [5] |
| | Prediction Block | 29.46 (0.43) | | | | 492–511 | |
| **Intact stimuli** | **Degraded stimuli** | | | | | 78–472 | |
| | Expected | 177.09 (2.19) | 78.1 | 136.7 | 58.04 | 500–523 | |
| Expected | Random Block | 43.92 (0.64) | | | | 531–558 | 113–222[29] |
| Random Block | | | | | | | 429–453[7] |
| | Expected | 177.09 (2.19) | 85.9 | 125.0 | 58.06 | 82–398 | 460–472[4] |
| | Prediction Block | 122.09 (0.79) | | | | | |
| **Intact stimuli** | **Degraded stimuli** | | | | | | |
| | Unexpected | 176.93 (2.73) | 82.0 | 132.8 | 55.95 | 85–429 | |
| Unexpected | Random Block | 44.34 (0.58) | | | | 437–449 | 128–152 [7] |
| Random Block | | | | | | | 171–222[14] |
| | Unexpected | 176.93 (2.73) | 85.9 | 125.0 | 56.63 | 85–218 | |
| | Prediction Block | 29.37 (0.48) | | | | | |
| **Intact &** | **No Prior Exposure** | | | | | | |
| **Degraded stimuli** | Predictors | 362.31 (3.07) | 335.9 | 335.9 | 51.40 | None | |
| | Random Block | 89.06 (0.69) | | | | | 117–132 [5] |
| Expected | | | | | | | |
| Prediction Block | Predictors | 362.31 (3.07) | 164.1 | 171.9 | 52.13 | 164–187 [7] | |
| | Prediction Block | 308.63 (1.04) | | | | 250–277 [8] | |
| **Intact &** | **Prior Exposure** | | | | | | |
| **Degraded stimuli** | Predictors | 361.99 (2.13) | 171.9 | 199.2 | 52.62 | 171–214 [12] | |
| | Random Block | 88.61 (0.59) | | | | | None |
| Expected | | | | | | | |
| Prediction Block | Predictors | 361.99 (2.13) | 168.0 | 191.4 | 52.71 | 168–218 [14] | |
| | Prediction Block | 308.75 (1.05) | | | | | |

*Table Supplement 3: Summary statistics for decoding analyses. First, expectancy conditions and sequence blocks from which data were obtained are reported. Note that Random Block stimuli occurred randomly with no structured predictive relationships. Classifiers trained to discriminate Random Block stimuli were cross-validated and tested on the same stimulus identity occurring at expected or unexpected sequence positions in the Prediction Block. Next, the average number of stimulus epochs contributing to each analysis are reported, with standard deviation in brackets. The top and bottom numbers report the training and testing data, respectively. Then, onset and peak times of decoding relative to stimulus onset (0 ms) are reported, along with peak accuracy in percentages. Finally, timepoints with at least moderate evidence for above-chance decoding and condition differences are reported, including cluster size in square brackets for condition differences. All reported timepoints survived cluster-based permutation tests. Data for the top panels are based on all participants (N = 40). Data for the bottom panel are split by prior exposure groups (n = 20 each).*
